# Microfluidics based exploration for quorum quenching genes in Antarctic microbiomes

**DOI:** 10.1101/2025.11.18.689007

**Authors:** Joseph White, Jorge Díaz-Rullo, Verónica Morgante, José Eduardo González-Pastor, Aurelio Hidalgo, Mercedes Sánchez-Costa

## Abstract

Quorum sensing (QS) is a form of microbial communication that enables each cell in a population to sense the total population density, so that gene expression can be modified accordingly. Quorum quenching (QQ) is the antagonistic disruption of this communication by competing organisms, with potential use in the ongoing human effort to control microbial populations. Previous studies have taken advantage of functional metagenomics to retrieve new QS/QQ genes, but the frequency of obtaining positive clones remained very low, suggesting the need for increased screening efficiency. Here, a new ultrahigh-throughput screening method was developed to search for genes encoding novel QQ genes based on functional metagenomics, microfluidics, and the development of an *Escherichia coli* reporter strain whose fluorescence is repressed in the presence of AHLs but restored by the expression of any gene that interferes with the QS system. This strain was transformed with a metagenomic short-insert library collected from Antarctic plant rhizospheres; an understudied extreme environment. The library was encapsulated in droplets containing single clones and sorted using fluorescence-activated cell sorting (FACS). After screening approximately 7,000,000 clones, around 200 were recovered and one positive hit was confirmed, showing a previously unreported mode of action that would have been difficult to detect using rational computational methods. These findings underline the potential of microfluidics to dramatically increase screening efficiency, while reducing costs and processing time, and they act as a proof of concept for the discovery of more genes involved in QS and other molecular mechanisms of interest in microbial ecology.

**IMPORTANCE:** Quorum sensing (QS) regulates essential microbial behaviours, including virulence, biofilm formation, and symbiotic interactions, making it a key target for ecological and applied microbiology. Disrupting QS through quorum quenching (QQ) has major potential in agriculture, medicine, and biotechnology as an alternative to antibiotics or chemical treatments. However, the discovery of QQ genes through brute-force approaches is limited by large screening efforts, resulting in high costs and long processing times. Our study introduces a microfluidics-based, ultrahigh-throughput functional screening platform that strongly improves screening efficiency. It allowed us to identify a novel QQ gene, predicted to be involved in a previously undescribed mechanism of QS disruption, from an under-explored extreme environment (Antarctic rhizosphere). These results demonstrate how microfluidics can optimise screening technologies, while unlocking the hidden functional diversity of microbial communities useful for biotechnological applications in microbial control.

## INTRODUCTION

Quorum sensing (QS) is a means of communication between microbial cells, allowing for cell density-dependent control of gene expression (1). In its most basic form, a chemical signal molecule, referred to as an ‘autoinducer’, is synthesised by microbial cells and released into the environment. The concentration of this autoinducer increases at roughly the same rate as the population density until a concentration threshold is crossed, whereupon the autoinducer activates a positive-feedback loop by binding to a transcriptional regulator which upregulates the expression of the autoinducer gene. Meanwhile, the same activated transcriptional regulator binds to other promoter regions to alter the phenotype of the community in response to its high cell density (1–3). This density-dependent communication enables a microbial species to build a cooperative response to, for example, improve defence or access to nutrients, or to differentiate into alternate morphological forms (4, 5).

Various types of autoinducer molecules have been described, but by far the most common are the acyl-homoserine lactones (AHLs), usually known as AI-1. They are particularly widespread in the Gram-negative proteobacteria (6, 7), while also having been detected in distantly related organisms, such as cyanobacteria and even archaea (8–10). A classic example of QS that uses an AHL as its autoinducer is found in the bacterium *Vibrio fischeri*, which provides the luminescence foor some marine animals. When this microorganism is distributed at low density in open water, their QS relay is inactive. However, once it inhabits the light-emitting organ of an animal, it quickly replicates within the confined space to a density where their AHL autoinducer reaches the threshold concentration required to activate their transcriptional regulator (LuxR), which then dimerizes and binds to specific promoters that contain a so-called ‘lux box’ to regulate their activity. In this way, LuxR regulates the expression of its own *luxR* gene and the lux operon, which contains the *luxI* autoinducer gene required for AHL biosynthesis, as well as the luciferase genes that provides bioluminescence. This positive autoregulation gives the system hysteresis, so that bioluminescence remains activated until the bacterial population density decreases again (1, 6, 11–17).

Important as it is in aiding the establishment and dispersal of microbial communities, QS has prompted the evolution of quorum quenching (QQ) systems that are able to disrupt it (18), allowing antagonist species to compete more effectively with the QS species in their environment. Mechanisms of QQ include inhibiting signal production, inactivating or degrading the signal, inhibiting the signal reception by competing with the signal molecule’s receptor analogues or by blocking the signal transduction cascade (19, 20). Such abilities have been observed in a huge variety of different organisms, from prokaryotes to fungi, animals and plants (21–26). There are three common classes of QQ enzymes that modify or degrade AHL autoinducers: lactonases, acylases, and oxidoreductases (27, 28). The diversity of such enzymes that has already been discovered suggests that these systems have evolved independently on several occasions, representing a clear example of convergent evolution (22, 29), further demonstrating the importance and widespread use of QQ systems in nature.

Considering that bacterial virulence is often triggered by QS systems, it is of little surprise that much interest has been directed towards QQ research, which represents a potential new means of defence against microbial threats, particularly in the light of increasing resistance against antimicrobial chemotherapy (30–32). Most of the QQ genes characterized so far have been discovered using traditional cultivation-based techniques. However, this is slow and laborious work, while also being untenable for the vast majority of prokaryotes, which remain unculturable (33). As such, more and more laboratories are turning to metagenomics to ‘mine’ environmental samples for genes of interest (34–36), but this brute-force approach also requires large screening efforts, resulting in high costs and long processing times, which significantly limit its chances of success. These are particularly relevant factors when the frequency of obtaining positive hits is very low as reported in previous functional metagenomics studies searching for QS genes (Torres et al., 2018). To address these challenges, microfluidics offers a powerful alternative enabling the manipulation of fluids at the micrometer scale by using picoliter-sized droplets as test-tube analogues. This approach offers several advantages: i) reduced volume, leading to lower reagent consumption and costs; ii) increased throughput; iii) signal concentration within a confined space, and iv) minimized plastic use (37–41). Therefore, in this work, a fluorescent QQ reporter strain, developed using modified genetic components taken from the *lux* operon of *V. fischeri*, was used in conjunction with cultivation-independent functional metagenomics and high-throughput microfluidic screening coupled to fluorescence-activated cell sorting (FACS) to search for novel QQ genes from microorganisms harvested from Antarctic rhizospheres.

## EXPERIMENTAL PROCEDURES

### Bacterial strains, enzymes, oligonucleotides, constructs, media and culture conditions

*Escherichia coli* DH10B (Invitrogen) was routinely grown in Lysogeny broth (LB) medium at 37 °C, 30 °C or 26 °C, depending on the type of plasmid it contained and the experiment. Cultures were incubated routinely at 200 rpm, or 50 rpm when incubated on a Fisherbrand 3D Platform Rotator (Fisher Scientific) to provide the swirling motion necessary to maintain 3-oxo-C10-HSL suspension. When required, media was supplemented with 10 μM of 3-oxo-C10-HSL. If appropriate, media were supplemented with sterile-filtered solutions of antibiotics tetracycline (TET) 10 μg ml^−1^, when selecting for clones transformed with the λ-red plasmid; chloramphenicol (CHL) 40 μg ml^−1^, when selecting for clones successfully recombined with the *cat_luxR_gfp* cassette; and ampicillin (AMP) 100 μg ml^−1^ for *E. coli* strains transformed with pBluescript II SK + (pSKII+). Restriction enzymes, polymerases and T4 DNA ligase were from New England Biolabs (UK) or Roche (Germany). Other chemicals were from Sigma-Aldrich (Germany) and were at least of reagent grade purity. Electroporation was carried out using a Micropulser (Bio-Rad) according to the manufacturer’s instructions.

### Construction of DH10B Δ*rbsAR*::*cat_luxR_gfp* reporter strain for quorum quenching

The quorum quenching reporter cassette *cat_luxR_gfp* was constructed in three parts chemically synthesized by IDT (Coralville, IA, USA). First, the CHL resistance *cat* gene with Shine-Dalgarno sequence (BioBrick BBa_J31005) was combined with an upstream promoter sequence (BioBrick BBa_I14033) and downstream sequence T10 terminator taken from the pSB1C3 plasmid, as found on the iGem parts registry (http://parts.igem.org/Main_Page), with non-coding sequences maintained (except for the addition of a *BamHI* restriction site downstream of the promoter). Second, the *luxR* gene from *V. fischeri* (BioBrick BBa_C0062) was fused with a promoter (BBa_J23119) reverse orientated to avoid read-through, with three base changes at positions 54T>A, 174C>A and 627C>T to remove restriction sites *HindIII*, *XbaI* and *PstI*, and with an artificial Shine-Dalgarno sequence, designed using the Salislab website (https://salislab.net/software/OperonCalculator_EvaluateAnnotatedOperon) (42). The *luxR* terminator was the modified *thr* terminator (BBa_B1006). Third, the *gfp* gene was taken from pET28b.sfGFP together with its Shine-Dalgarno sequence (43), and expressed under a modified version of the truncated Pλphage constitutive promoter used by Pinheiro and collaborators (44). The ‘lux box’ sequence was inserted between the −10 and −35 regions of the truncated Pλphage promoter, so that LuxR functioned as an AHL-dependent repressor at this promoter, rather than as a transcriptional inducer as in the AHL autoinducer system of *V. fischeri* (Fig. 1) (45).

**Figure 1.**
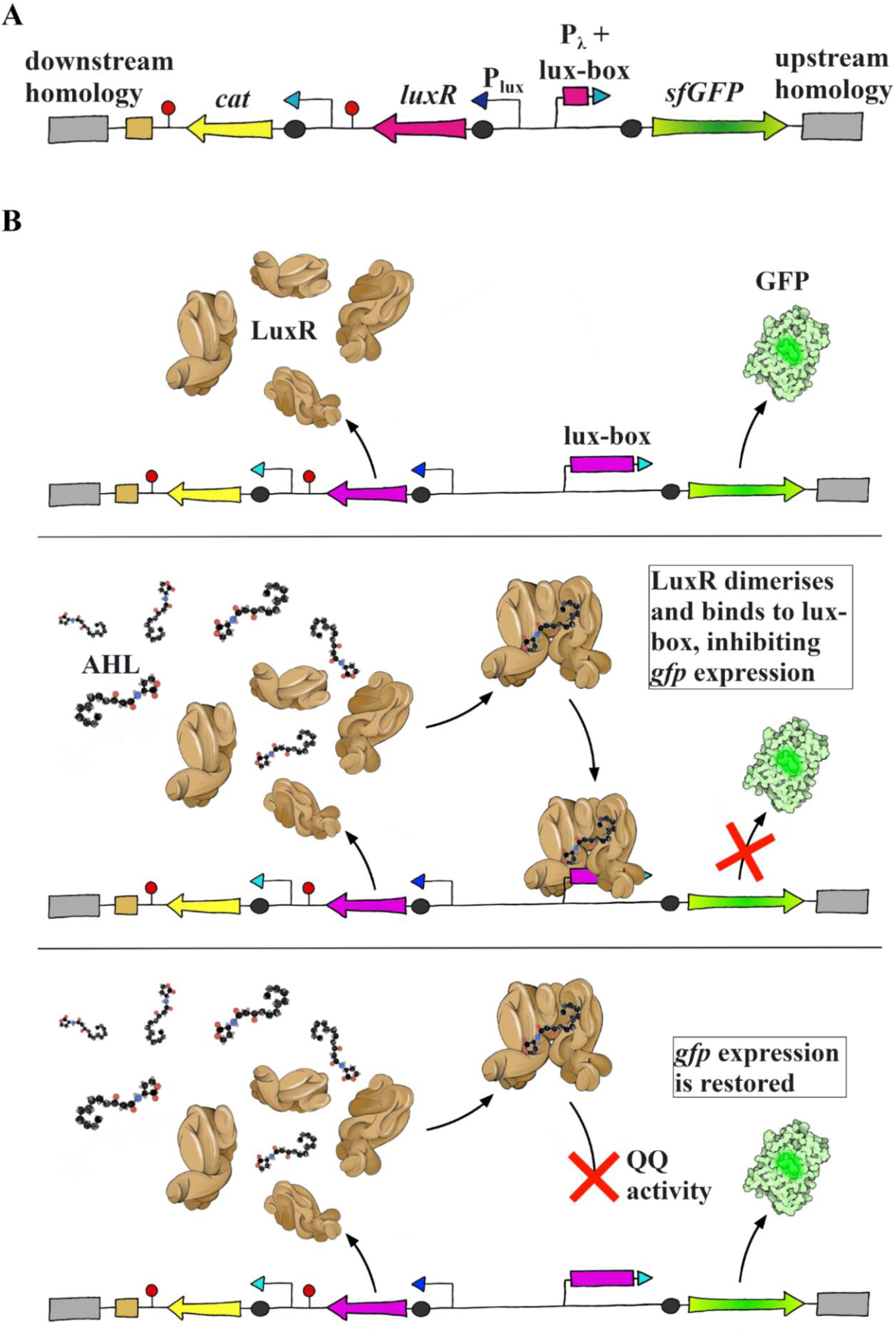
Design and function of the *cat_luxR_gfp* reporter cassette. (A) Genetic construct integrated at *rbsAR* locus. *cat*: chloramphenicol acetyltransferase gene; *luxR*: luxR family transcriptional regulator gene; Pλ+lux box: truncated constitutive Pλphage promoter with luxR binding site, causing repression in the presence of luxR + homoserine lactone; sf-gfp: super-folding green fluorescent protein gene; black circles: RBS sites; red lollipops: terminator regions. (B) Top: The *luxR* gene produces LuxR protein under constitutive expression and GFP under expression of the Pλphage promoter with lux-box. Center: The LuxR protein binds to the AHL present in the medium. Once bound to AHL, the LuxR protein dimerizes and is able to bind to the lux-box region of the truncated Pλphage promoter sequence, upstream of the *gfp* gene. In doing so, it represses expression, and the GFP phenotype is abolished. Bottom: If a QQ system is in play, the AHL will be made unavailable or the luxR will be prohibited from binding to the DNA. Either way, *gfp* expression will remain active.

The three synthetic parts of the whole construct were assembled by restriction, digestion and ligation. Subsequently, they were integrated into the *rbsAR, arsB and lacZ* loci of the *E. coli* DH10B chromosome by linear DNA recombination using the λ-red system (46). To facilitate the homologous recombination, 200 bp homology arms were introduced at 5’-end of the *cat* cassette and at the 3’ end of the *gfp* cassette. Competent *E. coli* DH10B cells were first transformed with the λ-red plasmid by electroporation and the transformation plates incubated in the dark at 30 °C overnight. The next day, LB-TET was inoculated with single colonies from the transformation and incubated at 30 °C in the dark, overnight. LB-TET supplemented with 0.35 % L-arabinose was then inoculated with the previous day’s overnight cultures (1:1000 dilution) and incubated in the dark at 30 °C until the OD_600_ reached 0.2. At this point, they were grown at 37 °C to induce the recombinant proteins of the λ-red plasmid, while also inhibiting plasmid replication, and incubated until the OD_600_ reached 0.4. Competent λ-red DH10B *E. coli* cells were transformed with 1-2 μg of the linear DNA cassette. Directly after transformation, the cells were re-suspended in 1 ml of S.O.C. medium (ThermoFisher Scientific) supplemented with 0.3 – 0.4 % L-arabinose and incubated at 37 °C overnight. The next day, these cultures were spread onto LB-CHL agar plates and incubated at 37 °C to select for recombinant clones. Modified regions were sequence verified.

To test the functionality of the reporter strain, the quorum-quenching *hqiA* gene, encoding a lactonase (47), was excised from the original pME6010 plasmid by digestion with *BglII* and *HindIII*. The resulting fragment was ligated into the multicloning site of pSKII+, previously digested with *HindIII* and *BamHI*. The resulting construction (pSKII+-*hqiA*) was electroporated into the reporter strain previously described. A strain transformed with an empty pSKII+ plasmid was used as the negative control. Both strains were grown in LB with and without 3-oxo-C10-HSL at 26 °C for 48 hours. Cultures were diluted in phosphate buffered saline (PBS) to an OD_600_ of 1, and fluorescence measurements were performed using a Qubit™ 3 Fluorometer (Invitrogen) (λ_ex_ 430 to 490 nm, λ_em_ 510 to 580 nm). All measurements were performed in triplicate. Microscopy was carried out with a fluorescence microscope Leica CTR6000 (Leica Microsystems) coupled with a digital camera Hamamatsu C11440 (Hamamatsu Photonics Deutschland GmbH) using an excitation at 500/24 nm and an emission at 542/27 nm when capturing fluorescence images.

### Development of an ultrahigh-throughput screening assay

The fabrication of polydimethylsiloxane (PDMS) chips was carried out as previously described (48, 49). Sylgard 184 silicone elastomer and Sylgard 184 curing agent (Sylgard) were mixed at a mass ratio of 10:1, degassed in a liophilizer, poured on a clean silanized master photolithographic mask with the embossed design for a 2-inlet, 20 μm flow-focussing device (available at https://openwetware.org/wiki/DropBase), and degassed a second time. The covered masters were cured at 60 – 70 °C for > 3 hours, PDMS slabs were carefully cut away and holes were punched for the inlets/outlets with a 1 mm diameter retractable biopsy punch (KAI Medical). Chips were washed by immersion in a bath of isopropanol with ultrasound for 15 min, then washed in a bath of water with ultrasound for 15 min, before a final clean with nitrogen gas and Scotch tape. Chips and clean glass slides were then treated with oxygen plasma (200 W, 150 ml/min) and joined. To make fluorophilic chips, channels were flooded with a 1 % solution of trichloro(1H,1H,2H,2H-perfluorooctyl) silane in HFE7500 (3 M) fluorinated oil and incubated overnight at 65 °C. To make hydrophilic chips, channels were flooded with 2 mg/ml poly(diallyldimethylammonium chloride) in 0.5 M NaCl, incubated at room temperature for 10 min, flooded with 150 mM NaCl, then with 2 mg/ml poly(sodium 4-styrenesulfonate) and incubated at room temperature for a further 10 min. Finally, the chips were flooded with ultrapure water and stored in moist paper in a sealed container at room temperature. All solutions were filtered through a 0.22 μm filter before use.

Cultures were grown overnight in LB-AMP-3-oxo-C10-HSL at 26 °C with swirling agitation, then re-suspended in quarter-strength LB-AMP-3-oxo-C10-HSL to an OD_600_ of 0.02. Single-cell encapsulation was achieved by injecting the diluted cell culture and a 0.45 μm filtered solution of 1.5 % (w/w) Fluosurf (Emulseo, France) in HFE7500 containing 10 μM 3-oxo-C10-HSL through a fluorophilic flow focussing chip at flow rates of 100 and 1000 μL·h^-1^, respectively. At this culture dilution, the encapsulation of one cell per droplet in ca. 9.5 % of the droplets was achieved, according to the Poisson distribution. Droplets, each around 4.5 pL, were collected in a low binding 1.5 ml tube, the emulsion was spun briefly and the excess oil removed with a syringe. Droplets were visualized in Fast-Read 102 slides with counting chambers (Biosigma s.r.l., Italy) using an Olympus BX50 microscope equipped with FITC fluorescence filters and a Pike F-032B camera (Allied Vision Technologies) and a 25X objective to confirm the droplet occupancy and integrity before being incubated at 26 °C overnight, with gentle agitation. After 12 h of incubation, the droplets were again imaged to confirm cell growth within the occupied droplets. Droplets were then re-encapsulated in 1 % (v/v) PBS-Tween using a 20 μm, hydrophilic, flow-focussing chips with a PBS:emulsion flow-rate of 100:10. All solutions and emulsions were driven through the chips using the Cetoni Nemesys Base 120 module and the Nemesys S pumps (Cetoni GmbH) and SGE gastight syringes of adequate volume connected to microbore polyethylene tubing (Smiths Medical). The microfluidic equipment was integrated by an inverted microscope (Leica DMi8) connected to a high-speed camera (Fastcam Mini UX 50, Photron) for real-time visualization of encapsulation experiments.

Emulsions were analysed and sorted using a FACSVantage SE (Becton Dickinson) at approximately 400 events per second. 44,100 and 50,595 events were analysed for the positive and negative control emulsions, respectively. Forward-scatter height (FSC-H) versus fluorescence plot was used to differentiate between fluorescent versus non-fluorescent droplets.

### Construction and screening of a metagenomic library from Antarctic plant rhizospheres

The fluorescent reporter strain DH10B Δ*rbsAR*::*cat_luxR_gfp* was transformed with a metagenomic library constructed with environmental DNA obtained from the rhizospheres of two Antarctic plants: *Colobanthus quitensis* (Antarctic pearlwort) and *Deschampsia Antarctica* (Antarctic hair grass), the only two flowering plants native to Antarctica. This original combined library contained around 1,000,000 clones with insert sizes ranging from 2 – 5 kb (50). DH10B Δ*rbsAR*::*cat_luxR_gfp* was transformed with a pool of plasmids isolated from the original library, then grown in LB-AMP-3-oxo-C10-HSL until the cell culture reached an OD_600_ = 10. The amplified new library was kept with 10 % glycerol (v/v) at –80 °C.

To screen the library, a 0.5 ml-aliquot of the library was thawed and used to inoculate 1 ml of LB-AMP-3-oxo-C10-HSL. Cultures were grown overnight at 26 °C with swirling agitation, then re-suspended in quarter-strength LB-AMP-3-oxo-C10-HSL to an OD_600_ of 0.02 and encapsulated, incubated and reemulsified as described above for the control populations. Emulsions were sorted by FACS using the previously established gating. Positive hits recovered from FACS were suspended in 1 ml PBS and spread directly onto LB-AMP agar plates. After incubation, recombinant colonies were replicated onto fresh plates and used to inoculate liquid LB-AMP supplemented with 3-oxo-C10-HSL. After 48 h incubation at 26 °C, all cultures were tested for green fluorescence using a benchtop Qubit™ 3 Fluorometer (Invitrogen). Clones testing positive under these conditions were re-streaked twice more, to ensure pure clonal colonies before being grown in culture. Plasmids were isolated from these cultures using a QIAprep Spin Miniprep Kit (QIAGEN, Hilden, Germany) before being used to transform DH10B Δ*rbsAR*::*cat_luxR_gfp*, to rule out false positives caused by spontaneous chromosomal mutations. New transformants were re-streaked twice before being used to inoculate LB with and without 3-oxo-C10-HSL, alongside the original recovered hits and the positive and negative controls, and incubated at 26 °C for 48 hours. Resultant colonies were streaked twice to again ensure pure clonal colonies before final cultures were incubated and tested for fluorescence as before.

### Characterization of hits

The positive hit recovered from FACS and confirmed by fluorescence testing after re-transformation was sequenced by primer walking, and putative ORFs were predicted using ORF Finder (https://www.ncbi.nlm.nih.gov/orffinder/) available at the NCBI website. For translation of the predicted ORFs, bacterial and archaeal codes were selected, allowing for the presence of alternative start codons. Predicted ORFs longer than 75 bp were used as queries in protein-protein BLAST searches against non-redundant protein sequences for functional assignment. Two halves of the original insert harbouring the putative identified ORFs were cloned out of the plasmid by PCR using the plasmids reported in Table S1, and inserted into empty pSKII+ by standard digestion and ligation with *Xbal*/*BamHI* (pAnt1-*orf*1) and *HindIII*/*BamHI* (pAnt1-*orf*2/3) and fresh competent stocks of DH10B Δ*rbsAR*::*cat_luxR_gfp* were transformed with each construction. Obtained recombinant strains were confirmed by sequencing. Similarly, *ybbA* gene from *E. coli* was cloned in pSKII+ by PCR, digestion and ligation (Table S1). Newly transformed strains were re-streaked twice before being used to inoculate LB with and without 3-oxo-C10-HSL, alongside the original complete hit and the positive and negative controls and incubated at 26 °C for 48 hours. All cultures were then re-tested for fluorescence as described above in order to confirm which clone was responsible for the observed fluorescence of the hit. Additionally, proteins encoded by pAnt1-*orf*2/3 were submitted to AlphaFold2 for structure prediction and evaluation of the quality of the predicted folding (51).

### Testing for esterase/ABC-transporter activity

Cultures of DH10B Δ*rbsAR*::*cat_luxR_gfp* harboring empty pSKII+, pAnt1-*orf*2/3 or pSKII+-*hqiA* were grown overnight in quarter-strength LB at 37 °C 200 rpm before being adjusted to OD_600_ of 0.6. For preparing protein extracts, 1.5 ml of the OD_600_-adjusted cultures was centrifuged at 16,000 x g for 2 min and cell pellets were re-suspended in 300 µl of BugBuster® reactive (Novagen) by pipetting and vortexing for 10 s. After incubation at room temperature for 20 min, samples were centrifuged at 16,000 x g for 30 min at 4 °C, and supernatants containing the crude extracts were transferred to a clean tube. Fluorescence of each extract was measured in Qubit™ 3 Fluorometer (Invitrogen) for verifying that the same quantity of GFP was extracted from each strain. 10 µl of each extract was mixed with 100 µl of 0.1 × PBS with HSL at a final concentration of 1 mM. As a negative control, another reaction was performed without adding any protein extract. Reactions were incubated at room temperature for 1 h. 18 µl of reaction product was added to 2 ml tubes containing 582 µl of an overnight culture of the reporter strain DH10B Δ*rbsAR*::*cat_luxR_gfp* in quarter-strength LB and adjusted to an OD_600_ of 0.002. Cultures were incubated at 30 °C at 50 rpm for 16 h. As a positive control, 600 µl of the OD_600_-adjusted culture of the reporter strain was incubated without adding any reaction product. 600 µl of each overnight culture were centrifuged at 16,000 x g for 2 min and the cell pellet re-suspended in 200 µl PBS. Fluorescence was measured in in Qubit™ 3 Fluorometer (Invitrogen). Fluorescence values were corrected with the OD_600_ values of the final cultures and the fluorescence values of the protein extracts.

### Statistical analysis

Statistical parameters, including value of n, mean and standard deviation (S.D.), are reported in Figure legends. One-way ANOVA followed by Tukey post hoc analysis was used to calculate statistical significance using GraphPad Prism version 9.00 (GraphPad Software, La Jolla, CA, United States).

## RESULTS

### Construction of a reporter strain for quorum signal inhibition

A strain derived from *E. coli* DH10B that expressed a quorum-signal dependent fluorescence, designated as DH10B Δ*rbsAR*::*cat_luxR_gfp*, was designed and constructed to detect genes conferring quorum signal inhibition. The ‘fluorescence switch’ included a CHL resistance gene, a transcriptional regulator gene *luxR* under the control of a constitutive promoter, and a *gfp* gene under the control of a promoter containing a ‘lux box’ in a way that LuxR functioned as an AHL-dependent repressor at this promoter (45) (Fig. 1A) (see Methods). The rationale behind this QQ reporter strain is that presence of AHL would activate LuxR, repressing GFP expression, while if this quorum signal response is disabled by the activity of a QQ transgene, fluorescence would be restored (Fig. 1B). The reporter construct was inserted at *rbsAR, arsB and lacZ* loci of *E. coli* DH10B chromosome, since these genes were previously characterised as well non-essential insertion loci (52). However, only the strain in which the reporter construct was inserted at *rbsAR* operon showed a clear fluorescent signal.

### Functional validation of reporter strain

To test the functionality of this reporter construct, DH10B Δ*rbsAR::cat_luxR_gfp* strain was transformed with the gene *hqiA*, which encodes for a previously identified QQ lactonase and could therefore work as the positive control (DH10B Δ*rbsAR::cat_luxR_gfp*/pSKII+-*hqiA*). This lactonase, HqiA, shows homology with enzymes belonging to the cysteine hydrolase (CHase) group that have been shown to cleave ether, ester and amide bonds, and it has the ability to degrade a broad range of unsubstituted, oxo- and hydroxyl-substituted AHLs (29). The strain was also transformed with empty pSKII+ to obtain a negative control (DH10B Δ*rbsAR*::*cat_luxR_gfp*/pSKII+).

Because HqiA lactonase was previously reported to show significantly higher efficacy of degradation with long-chain AHLs such as 3-oxo-C10-HSL, and due to short-chain AHLs are more prone to temperature-dependent lactonolysis than long-chain AHLs, 3-oxo-C10-HSL was chosen for these functional assays (47, 53), being 10 µM 3-oxo-C10-HSL was the minimal concentration needed to abolish the signal (Fig. S1). Negative control showed strong fluorescence in standard LB media lacking AHL, but repressed fluorescence after growth in the presence of 3-oxo-C10-HSL, while the positive control strain maintained the same high level of fluorescence regardless of the 3-oxo-C10-HSL concentration of the medium (Fig. 2).

**Figure 2.**
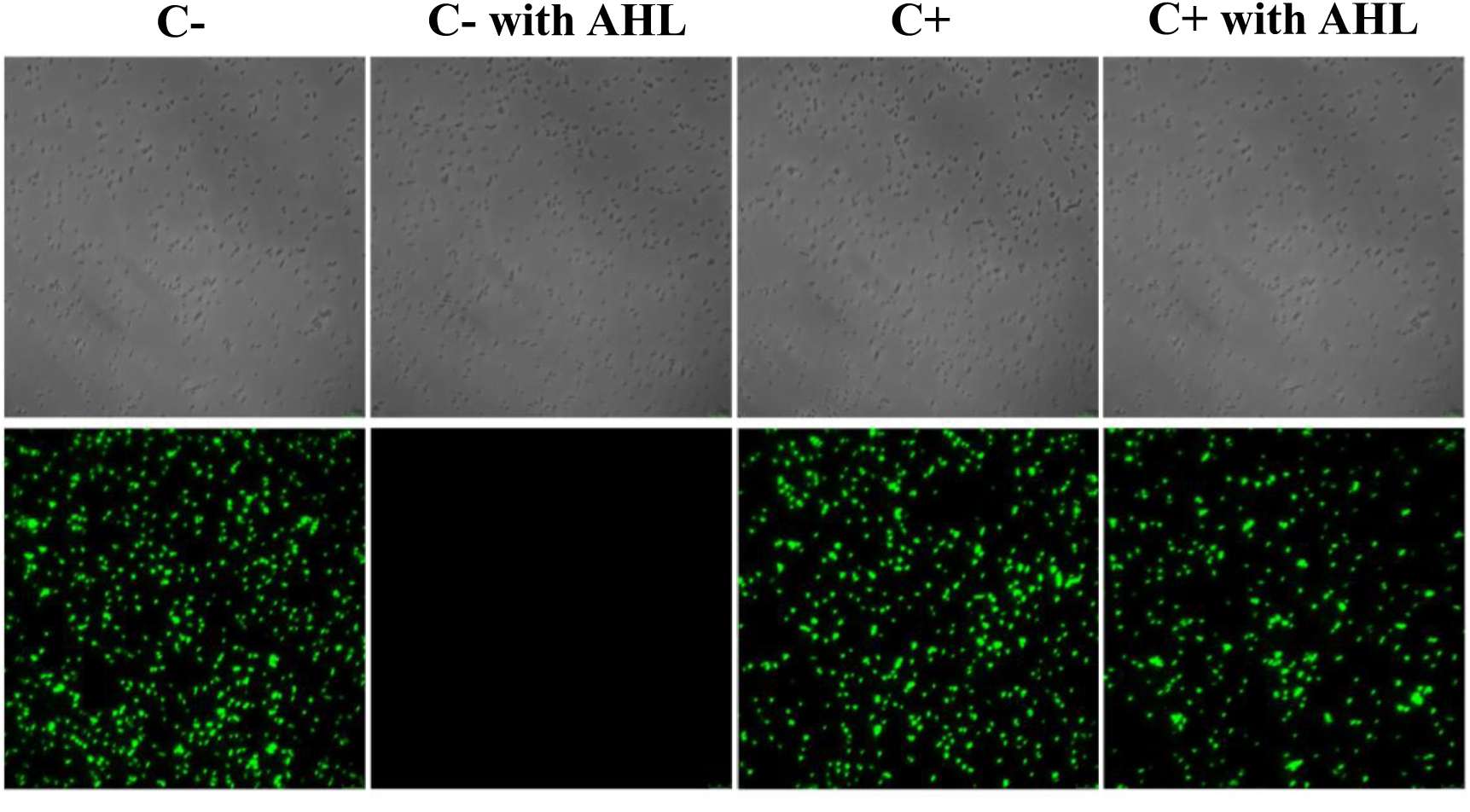
HqiA lactonase gene successfully re-establishes fluorescence in the presence of homoserine lactone. DH10B Δ*rbsAR*::*cat_luxR_gfp*/pSKII+ strain (negative control, C-) and DH10BΔ*rbsAR*::*cat_luxR_gfp*/pSKII+-*hqiA* strain (positive control, C+) were grown with or without 10 µM 3-oxo-C10-HSL (AHL). All cultures grown at 26 °C for 48 h before re-suspension in PBS to OD_600_ = 1. (top: phase contrast; bottom: green fluorescence; magnification: 1000X).

Considering the correct functionality of the reporter, we implemented this assay in water-oil-water (w/o/w) droplets using microfluidics. For that purpose, and prior to cell encapsulation, each control strain was suspended in LB diluted to 25 % (v/v). This step was necessary for reducing the background fluorescence produced by LB components that interferes with the signal picked up from a fluorescent-cell occupied droplet, while sufficient nutrients are still provided for overnight growth of encapsulated cells. Cells were double-encapsulated in w/o/w droplets to allow for cell sorting in the hydrophilic matrix of most commercially available FACS sorters. For double-encapsulation, cultures were first passed through a microfluidic chip to create aqueous droplets within a stable oil matrix, using a biocompatible oil phase that facilitates the transfer of respiratory gases (Fig. 3A). Cell-containing droplets were passed through a hydrophilic microfluidic chip, where each droplet was encapsulated again in an outer aqueous layer, creating double emulsions with an oil shell between two aqueous phases (54–57). Double-encapsulated cells were incubated at 26 °C overnight in the presence of 3-oxo-C10-HSL, as growth of the single cells would amplify the assay fluorescence signal and facilitate FACS sorting (Fig. 3B-C). This microfluidic approach enabled the determination of gating criteria for sorting positive events (Fig. 3C).

**Figure 3.**
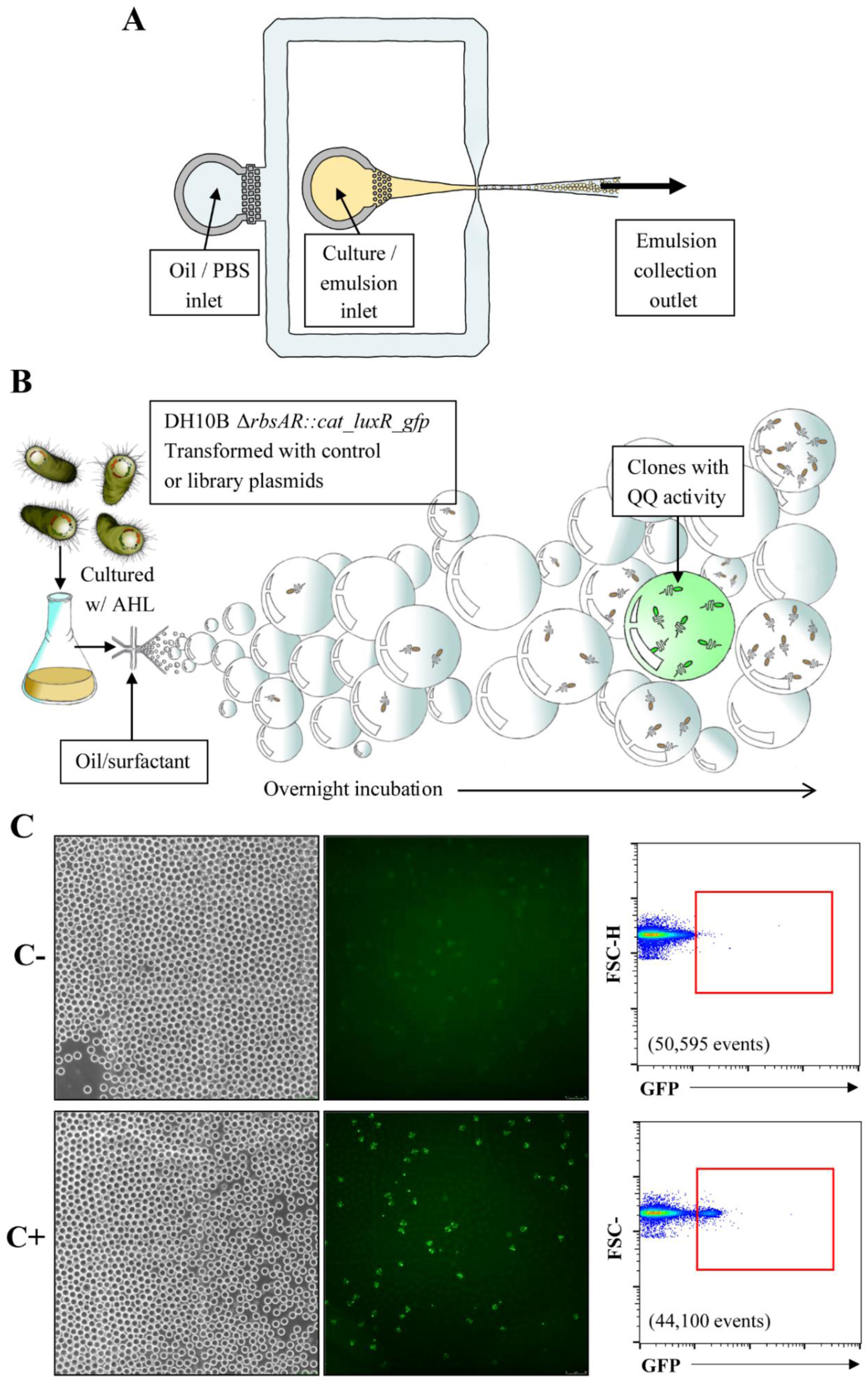
Microfluidic approach for sorting cells showing QQ phenotype. (A) Standard, single-intersection chip with 20 μM channel was used to encapsulate cells at a Poisson distribution to ensure a maximum of 1 cell per droplet (giving 10 % occupation). (B) Droplets harbouring DH10B Δ*rbsAR*::*cat*_*luxR*_*gfp*/pSKII+ (negative control, C-) or DH10BΔ*rbsAR*::*cat*_*luxR*_*gfp*/pSKII+-*hqiA* (positive control, C+) strains were incubated in the presence of 10 µM 3-oxo-C10-HSL at 26 °C overnight, allowing clonal cells to multiply within their droplets. Those expressing QQ activity also expressed green fluorescence, allowing detection by FACS. (C) Fluorescence microscope was used to take images of droplets harbouring control strains (left: phase contrast; right: green fluorescence; magnification: 20X). Droplets were sorted by FACS at approximately 400 events per second. 44,100 and 50,595 events were analysed for the positive and negative controls, respectively. Forward-scatter height (FSC-H) versus fluorescence plot was used to differentiate between fluorescent versus non-fluorescent droplets. Red boxes show population considered as positive.

### Screening of a metagenomic library from the rhizosphere of Antarctic plants to search for genes inhibiting QS

To identify genes involved in QQ, a functional metagenomics approach was performed using microfluidics to screen a large library from an understudied extreme environment. For that, the DH10B Δ*rbsAR*::*cat_luxR_gfp* reporter strain was used as the host of a previously obtained metagenomic library built in *E. coli* DH10B strain with the metagenomic DNA isolated from the rhizomes of the Antarctic vascular plants *C. quitensis* and *D. Antarctica* clones in pSKII+ plasmid (50). Single metagenomic clones were double-encapsulated in the presence of 3-oxo-C10-HSL and approximately 7,000,000 events were analysed and sorted as described above. Using gating parameters previously set using control strains, 400 positive hits were recovered from FACS (Fig. 4A) and plated on LB-AMP supplemented with 3-oxo-C10-HSL. After overnight incubation, we obtained 222 recombinant clones. QQ phenotype of these clones was validated by culture in the presence of 3-oxo-C10-HSL to fluorescence analysis. One recombinant clone, called pAnt1, was confirmed as positive (Fig. 4B), which corresponds to 0.5 % positive clones showing QQ phenotype among the 222 filtered recombinant clones. Moreover, considering that the sorted library contained 1,000,000 clones with a mean insert length between 2 and 5 kb, the frequency of positive clones obtained with this ultrahigh-throughput approach was 3 × 10^−4^ clones per Mbp.

**Figure 4.**
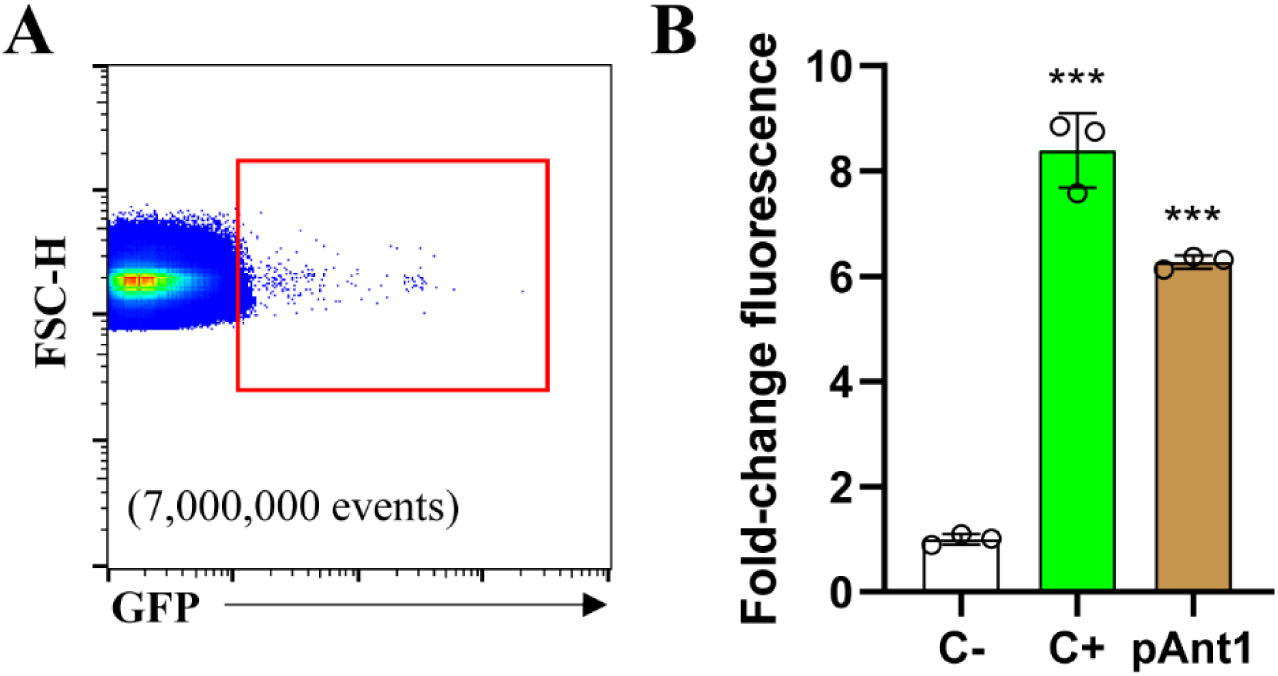
Sorting of Antarctic library and identification of positive clone pAnt1. **(A)** DH10B Δ*rbsAR*::*cat*_*luxR*_*gfp*/pAnt_library was double-encapsulated in droplets and incubated overnight at 26 °C in the presence of 10 µM 3-oxo-C10-HSL. Droplets were sorted by FACS at approximately 400 events per second, and approximately 7,000,000 events were analysed. Red box inside FACS plot shows population recovered using the same gating criteria fixed with the controls (Fig. 3). 400 positive hits were selected from FACS, and 222 recombinant clones were recovered. From these, one clone (pAnt1) showed a consistent quorum inhibiting phenotype. (B). Fluorescence of pAnt1 or DH10B Δ*rbsAR*::*cat*_*luxR*_*gfp*/pSKII+-*hqiA* (positive control, C+) in the presence of 10 µM 3-oxo-C10-HSL compared with DH10B Δ*rbsAR*::*cat*_*luxR*_*gfp*/pSKII+ (negative control, C-; fold-change). Strains were grown in 25 % LB supplemented with AMP and 3-oxo-C10-HSL for 48 h at 26 °C before resuspension in PBS to an OD_600_ of 1. One-way ANOVA followed by Tukey post hoc analysis revealed the displayed significant differences against the negative control (***p < 0.001). Data represent the mean ± S.D. (n = 3).

### Identification and characterization of the gene involved in QS inhibition

The complete sequence of the environmental DNA fragment harboured by the pAnt1 plasmid was obtained by primer walking. This fragment of 2 kb contained three putative ORFs likely to contain the gene of interest (Fig. 5). The pAnt1-*orf*1, located at the 5’-end of the fragment, encoded for a protein truncated at the C-terminal similar to a FtsX-like permease family protein from *Rhodanobacteraceae bacterium* (78.1% identity). pAnt1-*orf*2 and pAnt1-*orf*3 occupied a similar region on the insert, though in different orientations. The reverse-orientated pAnt1-*orf*2 encoded for a protein that matched with an ABC transporter ATP-binding protein, most similar to that found in the Gram-negative proteobacterium *Dokdonella fugitiva* (86.4% identity) and harbouring a conserved YbbA protein domain. Meanwhile, the C-terminal trunctated protein encoded by the forward-orientated pAnt1-*orf*3 showed similarity to an acyl-CoA thioesterase I precursor from *Leclercia adecarboxylata*, but only with a 32.7% identity.

**Figure 5.**
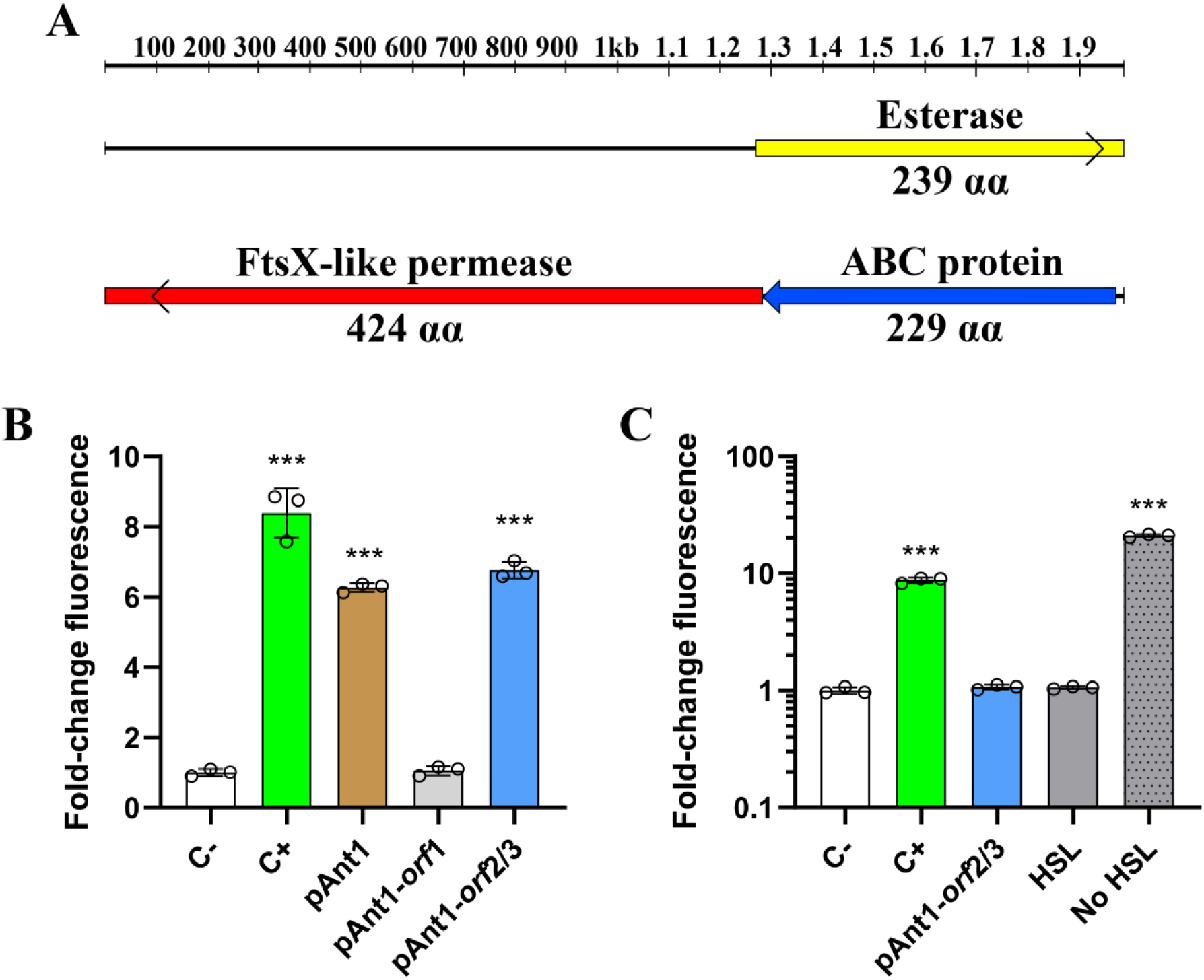
Characterization of pAnt1. (A) Schematic organization of the predicted ORFs identified in the fragment of eDNA harboured by the plasmid pAnt1. Arrows indicate the locations and the transcriptional orientation of the ORFs in the different plasmids. An open arrow near the corresponding end indicates truncated ORFs. (B) Fluorescence of DH10B Δ*rbsAR*::*cat*_*luxR*_*gfp* clones transformed with pSKII+ harboring *hqiA* lactonase gene (positive control, C+), pAnt1, pAnt1-*orf1* or pAnt1-*orf2/3* compared with reporter strain harboring empty pSKII+ (negative control, C-; fold-change). Strains were grown in 25 % LB supplemented with AMP and 10 µM 3-oxo-C10-HSL for 48 h at 26 °C before resuspension in PBS to an OD_600_ of 1. One-way ANOVA followed by Tukey post hoc analysis revealed the displayed significant differences against the negative control (***p < 0.001). Data represent the mean ± S.D. (n = 3). (C) Fluorescence of the reporter strain DH10B Δ*rbsAR*::*cat*_*lux*_*gfp* incubated with the products of the reactions of pure 3-oxo-C10-HSL with protein extracts obtained from DH10B Δ*rbsAR*::*cat*_*lux*_*gfp* harboring empty pSKII+ (negative control, C-), pSKII+-*hqiA* (positive control, C+) or pAnt1-*orf2/3*. Fluorescence was normalized with the measurements of negative control (C-; fold-change). 10 µL of protein extracts were mixed with 3-oxo-C10-HSL at a final concentration of 1 mM. After 1 h, reaction products were added to cultures of reporter strain, and cultures were incubated at 30 °C and 50 rpm for 16 h. Reporter strain was also incubated in the presence or absence of 10 µM 3-oxo-C10-HSL (HSL and no HSL conditions, respectively). One-way ANOVA followed by Tukey post hoc analysis revealed the displayed significant differences against the negative control (***p < 0.001). Data represent the mean ± S.D. (n = 3).

To determine which predicted ORF was responsible for the QQ phenotype, the upstream part of the plasmid containing ORF1 and encoding an FtsX-like permease family protein (pAnt1-*orf*1), and the downstream part harbouring ORF2 and ORF3 encoding the predicted esterase and the ABC transporter ATP-binding protein (pAnt1-*orf*2/3) was individually sub-cloned into the reporter strain. Subsequent fluorescence analysis confirmed all quorum-disrupting activity to be associated with the downstream predicted ORF region (pAnt1-*orf*2/3) (Fig. 5B). A possible explanation for this phenomenon could be that the predicted esterase activity of the protein encoded by pAnt1-*orf*3 degrades 3-oxo-C10-HSL, in a similar way to lactonases. The alternative hypothesis was that the ABC transporter ATP-binding protein encoded by pAnt1-*orf*2 may be part of an ABC transporter that could export 3-oxo-C10-HSL from the cell.

To address whether the positive phenotype exhibited by pAnt1-*orf*2/3 was caused by a lactonase-like activity that degraded 3-oxo-C10-HSL, protein extracts of the DH10B Δ*rbsAR*::*cat_luxR_gfp* strain harboring empty pSKII+, pAnt1-*orf*2/3 or pSKII+-*hqiA* were obtained and incubated with 3-oxo-C10-HSL. The reaction products were then added to cultures of the reporter strain and fluorescence was measured. As expected, an increase in fluorescence values was observed when the reporter strain was incubated with the protein extract obtained from the positive control pSKII+-*hqiA* strain expressing the HqiA lactonase (Fig. 5C). However, no differences in fluorescence values were observed when the reporter strain was incubated with the protein extract obtained from the strain harboring pAnt1-*orf*2/3 (Fig. 5C). Instead, it showed similar fluorescence activity to that incubated with the empty pSKII+ (negative control) protein extract (Fig. 6C). This result suggests that the pAnt1-*orf*2/3 QQ phenotype is not related to an AHL-degrading process. In addition to that, the AlphaFold2-predicted structure of the proteins encoded by pAnt1-*orf*2/3 exhibited low confidence metrics for the putative esterase, with a predicted local distance difference test (pLDDT) score below 50 and substantial regions of disordered conformation (Fig. S2A).

In contrast, folding simulations of the predicted ABC transporter ATP-binding protein showed a high confidence (pLDDT > 90) (Fig. S2B). Additionally, the uncharacterized ABC transporter ATP-binding protein YbbA protein from *E. coli* was the closest match to the putative ABC transporter ATP-binding protein encoded by ORF2 out of the structurally characterized proteins available in UniProt/Swiss-Prot (52% identity). As such, it seemed a more likely candidate, so we over-expressed YbbA in the reporter strain. However, the QQ phenotype was still absent (Fig. S3). These data suggest that pAnt1-*orf*2 would encode for a protein showing a new biological activity related to QS inhibition.

## DISCUSSION

With the creation of a new QQ reporter strain and its application in a microfluidics-based high-throughput screening of a metagenomic library, this work has detected a novel QQ gene from a hitherto little-investigated environment (Antarctic rhizospheres) and in doing so also provided proof-of-principle for the viability of this approach.

Transgenes responsible for creating the fluorescence switch of the reporter strain included a transcriptional regulator gene *luxR*, under the control of a constitutive promoter, as well as a super-folding green fluorescent protein (*gfp*) gene under the control of a promoter with a ‘lux box’ – a highly conserved, LuxR-binding DNA sequence – situated between its −10 and −35 sites. Genetic elements were recombined onto the chromosome of the reporter strain, rather than being expressed from a plasmid, to increase stability and give a more consistent GFP signal without the added variability of plasmid copy number. For that purpose, insertional locus of the reporter construct in *E. coli* genome was critical, since an efficient chromosomal integration of large DNA fragments tends to be problematic. Furthermore, both the *luxR* transcriptional regulator gene and the ‘lux box’ sequence from the *lux* operon of *V. fischeri* were modified in this work, so that the *luxI* (autoinducer) gene was absent, the *luxR* gene was constitutively expressed, and only the *gfp* transgene was under the control of a ‘lux box’ promoter. Normally, LuxR protein behaves as a transcriptional activator, but in this instance the ‘lux box’ was modified to repress rather than amplify transcription with LuxR binding (58), which is a relatively simple alteration, since the position of the lux-box with respect to the promoter is thought to help determine its strength and function as an activator or a repressor. Although this logic response has been questioned (59), context is so crucial in promoter function that it continues to elude predictive computation (60, 61), and the AHL-repressible promoter designed by Egland and Greenberg has since undergone further validation (62). The expression-repressing lux-box sequence and position described by Egland & Greenberg was tested in this design and, upon showing strong LuxR-controlled repression of the *gfp* gene, was selected to work with (45).

This new QQ reporter offers some advantages over previous tools used in metagenomic studies for the identification of new QS/QQ genes, such as the fluorescent *E. coli* AHL reporter, that emits fluorescent signal in the presence of AHLs (63), or the *Chromobacterium violaceum* CV026 biosensor, bacteria that produce a violaceus compound when exposed to AHLs (47, 64, 65). Among them: i) our reporter strain is designed to express fluorescence only when any QQ gene is present in the cell; ii) it is sensitive and at least semi-quantitative, with variations in QQ activity leading to different levels of GFP expression, and iii) it would be sensitive to any gene product that interferes with the QS system, rather than being calibrated to identify enzymes that degrade particular forms of AHL (66).

Another crucial element that must be considered in these kinds of studies is the AHL chosen. All AHLs are composed of a homoserine lactone ring connected via an amide bond to an acyl fatty acid side chain. They tend to vary in their composition by the length of this side chain (generally 4 - 18 C atoms), the presence of unsaturated bonds within it and the existence, or not, of an oxo- or hydroxyl group at the 3′ position (6, 67, 68). In all their forms, AHL molecules undergo lactonolysis, that is, the opening of the lactone ring by addition of a water molecule in aqueous solutions. However, this process is slowed by a reduction in temperature and pH, while AHLs with longer acyl chains are also naturally more resistant (69–74). In addition, it was reported that the positive control HqiA lactonase exhibited a higher esterase activity with long-chain AHLs such as 3-oxo-C10-HSL compared to AHLs with shorter chains. For these reasons, 3-oxo-C10-HSL was chosen in this work, allowing for 48 h of cultivation in swirling motion without any loss of signal, while abolishing fluorescence signal at a concentration inside the biological range observed in nature (75, 76).

Furthermore, the identification of novel QS genes using functional metagenomics has been successfully addressed by several laboratories in the past (47, 63–65). Therefore, in this work, we have combined our reporter strain with functional metagenomics for the discovery of environmental QQ genes. A metagenomic library constructed from an understudied, extreme environment (the rhizome systems of two vascular Antarctic plants: *C. quitensis* (Antarctic pearlwort) and *D. Antarctica* (Antarctic hair grass), which are the only two flowering plants native to Antarctica) was used for the search. Better understanding of cold-adapted quorum systems could be valuable, particularly in medicine and industry (77). Moreover, the soil around rhizospheres is rich in QS-active microbes and is a hotspot for Gram-negative bacteria that produce AHLs. As a result, several soil bacterial genera have been found to produce enzymes that degrade AHLs (78–81).

In this work, the fact that only one QQ gene was discovered might suggest that while the experimental pipeline was functioning as expected, there was an apparent paucity of QQ genes within this Antarctic environment. However, it should be taken into account that the expected number of positive clones was very low, attending to data from previous screens for QQ genes (Table 1). It was estimated that the frequency of positive clones obtained using metagenome screening varied between 1 × 10^−4^ to 3 × 10^−3^ per Mbp (47, 63–65), which is very similar to the frequency reported here of 3 × 10^−4^ per Mbp. This suggests that the low abundance of QQ genes in the Antarctic soil rhizosphere is similar to that found in other non-extreme environments.

**Table 1.** Metadata from previous studies reporting the discovery of new QS/QQ genes through non-ultrahigh throughput screening.

| Ref. | Environment | Reporter | Screened clones | Insert length | Positive clones | Positive clones per screened Mbp | Positive clones per screened clones |
| --- | --- | --- | --- | --- | --- | --- | --- |
| Nasuno 2012 | Forest soil and an activated sludge from a coke plant in Japan | Fluorescent *E. coli* AHL reporter | 103,700 | 40,000 | 3 | 7·E-04 | 3·E-05 |
| Riaz 2008 | Montrond soil samples (France) | *C. violaceum* CVO26 biosensor | 10,121 | 40,000 | 1 | 3·E-03 | 9·E-05 |
| Tannières 2013 | Root microbiome of *S. tuberosum* plant treated with γ-caprolactone | *C. violaceum* CVO26 biosensor | 29,760 | 40,000 | 1 | 8·E-04 | 3·E-05 |
| Torres 2018 | Hypersaline soil from Spanish saltern | *C. violaceum* CVO26 biosensor | 250,000 | 40,000 | 1 | 1·E-04 | 4·E-06 |
| This study | Antartic plant root rhizosphere | Fluorescent *E. coli* QQ reporter | 1,000,000<br>(222 after FACS pre-enrichment) | 3,500 | 1 | 3·E-04<br>(after FACS pre-enrichment) | 5·E-03<br>(after FACS pre-enrichment) |

This work also took advantage of microfluidics to dramatically reduce time processing and costs. FACS pre-enrichment, analyzing 1,000,000 library clones through 7,000,000 droplet events, reduced manual validation to 222 candidates. This represents a 4,500-fold reduction in labor while maintaining detection sensitivity (3×10⁻⁴ per Mbp) comparable to other previous studies is which tradicional metagenomics was used to find novel genes involved in QS (Table 1). These data highlight the utility of high-throughput screening based on droplet sorting for saving time, cost and reagents that may not be commercially available (i.e., AHLs) and reducing the environmental footprint of the work when compared with plastic-based assays. Additionally, this technique can be performed using equipment available in multiple research facilities, contributing to high-throughput screening democratization.

The single hit that was recovered from the library proved to be a further surprise. Our data suggests that the protein responsible for the QQ phenotype exhibited by pAnt1 is not related to lactonase AHL degradation, which differs from the novel QQ genes discovered in previous metagenomic studies (47, 63–65). In fact, pAnt1 analysis revealed that an ABC transporter ATP-binding protein might cause the reported QQ phenotype. To the best of the authors’ knowledge, no ABC-transporter has been previously shown to impart a QQ phenotype. Although it was largely assumed that all AHLs were freely diffusible in bacterial cells, such as in *V. fischeri* or *E. coli*, it is known that certain species, such as *Pseudomonas aeruginosa*, are not permeable to certain AHLs. In this bacterial species, the *mexAB-oprM* efflux pump is involved in the active efflux of both long-chain AHLs like 3-oxo-C10-HSL, and short-chain AHLs like C4-HSL, and the inactivation of this pump leads to the intracellular accumulation of these AHLs (82). Therefore, one explanation for these data is that the predicted ABC transporter ATP-binding protein encoded by pAnt1 has been retrieved from some environmental microorganism that uses active transport through ABC transporters to export AHLs out of the cell. However, if the observed QQ phenotype was caused by AHL efflux, the environmental ABC transporter ATP-binding protein should be able to interact with any *E. coli* ABC transporter permease with AHL transport potential, and to our knowledge, no AHL active transporter has been reported to date. Thus, deeper characterization of this ABC protein will be required to understand whether it represents a new QQ system that works by AHL active transport. The novelty of this putative QQ gene underlines the importance of using screening approaches not based only on sequence analysis, such as functional metagenomics, to discover new bacterial molecular mechanisms.

In conclusion, we have developed the first ultrahigh-throughput functional screening for the discovery of new QQ genes based on: i) microfluidic droplet sorting; ii) development of a fluorescent QQ reporter strain, iii) construction of a metagenomic library from Antarctic plants rhizosphere. This work verifies the importance of microfluidics for strongly increasing efficiency and reducing costs when performing functional screenings, and provides a novel fluorescent reporter strain sensitive to any gene product that interferes with the QS system. Furthermore, this naïve strategy allowed for the discovery of a novel QQ gene with a potentially previously unreported mode of action that would have been difficult to detect using computational methods. Libraries from other environments, large-insert libraries or different reporter strains based on other molecular mechanisms will be used in future ultrahigh-throughput screenings for expanding our knowledge of quorum systems.

## DATA AVAILABILITY STATEMENT

The datasets presented in this study can be found in online repositories. Names of the repository/repositories and accession number(s) can be found below: https://www.ncbi.nlm.nih.gov/nuccore/PP836139

## AUTHOR CONTRIBUTIONS

Experiment design and performance: J.W., J.D.R., V.M., A.H., J.E.G.P., M.S.C.; data analysis: J.W., J.D.R., M.S.C.; manuscript writing and reviewing: J.W., J.D.R., V.M., A.H., J.E.G.P., M.S.C.

## ACKNOWLEDGMENTS

We thank Inmaculada Llamas (Institute of Biotechnology, Biomedical Research Center (CIBM), University of Granada, Granada, Spain) for the kind gift ofpME6010::*hqiA* plasmid. We also thank Carolina González de Figueras from Centro de Astrobiología (INTA-CSIC). Finally, we aknowledge the Metafluidics project (Grant Agreement no. 685474) within the framework of the Horizon 2020 programme for its contribution to the conceptualization and early development of this work.

