## Supplemental figures S1, S2 and S3, and supplemental Table S1 for "Microfluidics based exploration for quorum quenching genes in Antarctic microbiomes"

[1]Department of Molecular Evolution. Centro de Astrobiología (CAB), CSIC-INTA. Carretera de Ajalvir km 4, Torrejón de Ardoz (28850). Madrid, Spain.

[2]Department of Molecular Biology. Universidad Autónoma de Madrid. Campus de Cantoblanco, Madrid 28049, Spain.

[3]Centro de Biología Molecular Severo Ochoa (UAM-CSIC) Nicolás Cabrera 1, Madrid 28049, Spain.

[4]Instituto de Biología Molecular. Universidad Autónoma de Madrid, Nicolás Cabrera 1, Madrid 28049, Spain.

[#] indicates co-first authorship. Author order was determined by drawing straws.

* Corresponding authors: Mercedes Sánchez-Costa and Aurelio Hidalgo

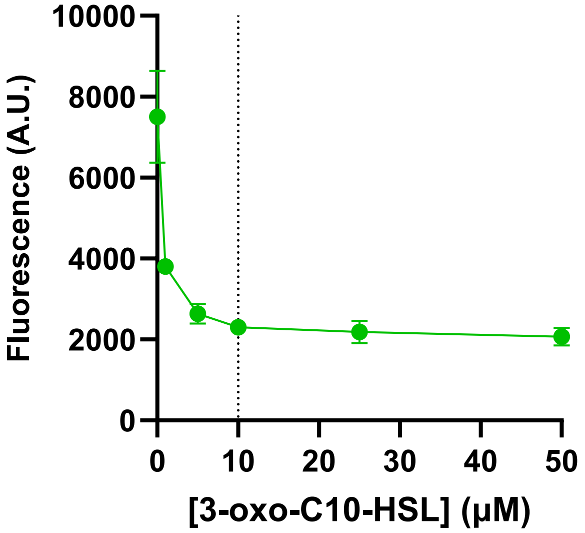


**Figure S1. Fluorescence signal of DH10B Δ*rbsAR*::*cat_luxR_gfp* reporter strain in the presence of different 3-oxo-C10-HSL concentrations.** Reporter strain was grown in 25 % LB supplemented with 3-oxo-C10-HSL for 48 h at 26 °C before resuspension in PBS to an $OD_{600}$ of 1. Vertical line indicates the selected working concentration of 3-oxo-C10-HSL.

**A**

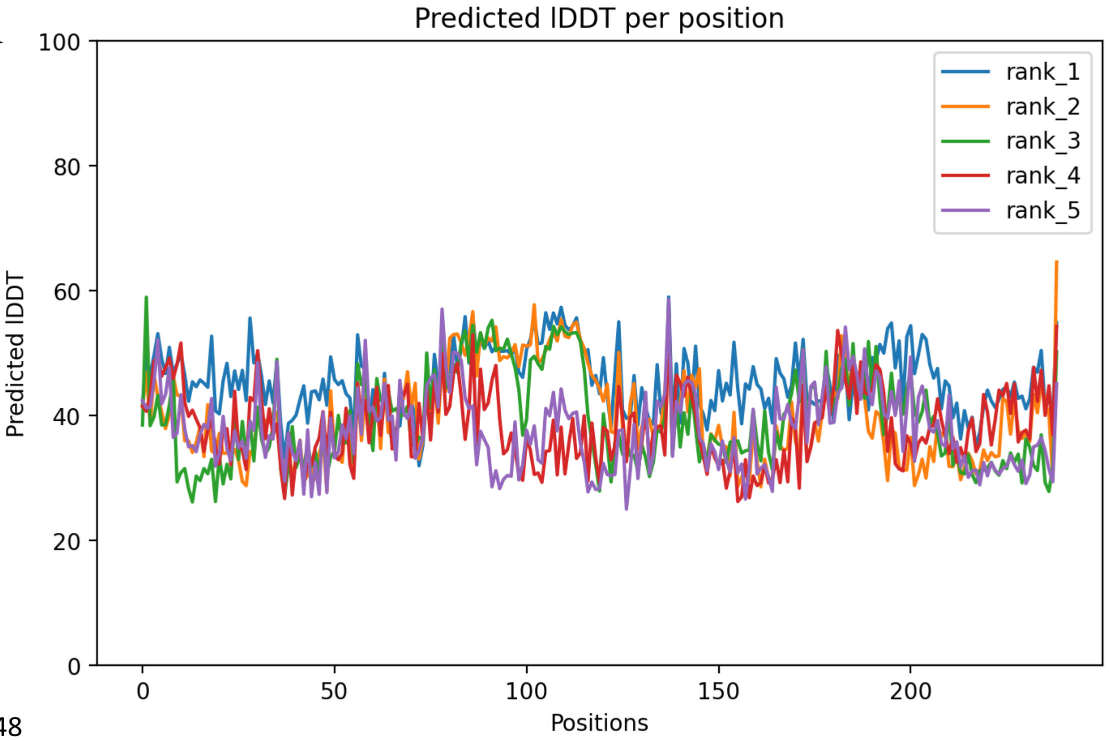


**B**

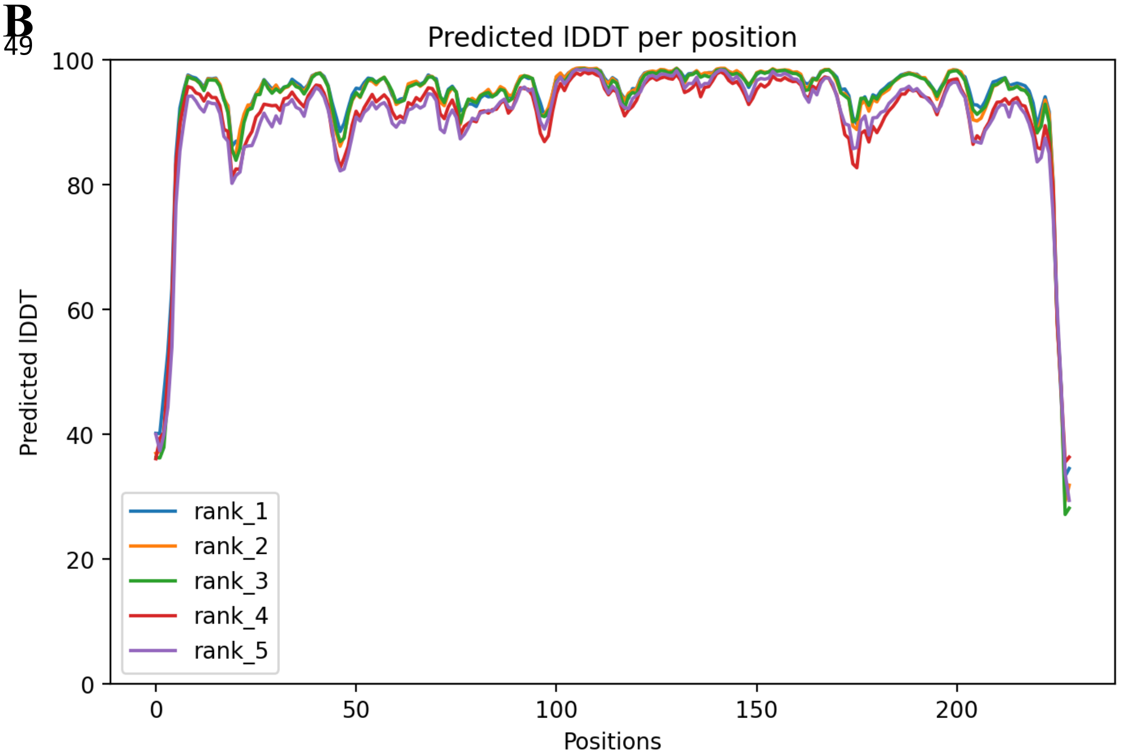


**Figure S2. Predicted local distance difference test (pLDDT) score plots of AlphaFold2-predicted structure of proteins encoded by pAnt1-*orf*2/3.** (A) Data from putative esterase encoded by pAnt1-*orf*3. (B) Data from putative ABC transporter ATP-binding protein encoded by pAnt1-*orf*2.

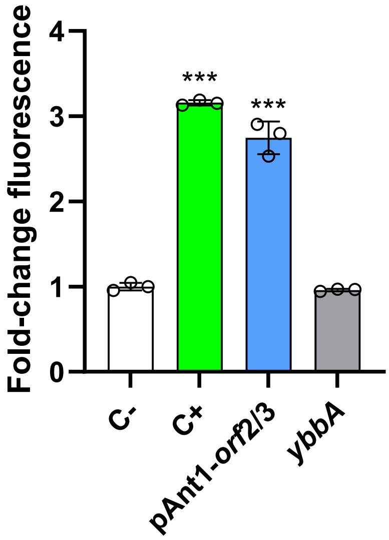


**Figure S3. *ybbA* gene from *E. coli* did not exhibit a QQ phenotype.** Fluorescence of DH10B Δ*rbsAR*::*cat_luxR_gfp* clones transformed with pSKII+ harbouring *hqiA* lactonase gene (positive control, C+), pAnt1-*orf2/3* or pSKII+-*ybbA* compared with the fluorescence of the reporter strain transformed with empty pSKII+ (negative control, C-; fold-change). Strains were grown in 25 % LB supplemented with AMP and 10 µM 3-oxo-C10-HSL for 48 h at 26 °C before resuspension in PBS to an $OD_{600}$ of 1. One-way ANOVA followed by Tukey post hoc analysis revealed the displayed significant differences against the negative control (***$p < 0.001$). Data represent the mean ± S.D. ($n = 3$).

**Table S1. Primers used to clone the *orf*1 and *orf*2/3 sections from pAnt1.**

| **Name** | **Sequence (5' → 3')** |
|---|---|
| pAnt_*orf1*_F | tagtacTCTAGACTCGAAGGACGCATTGAC |
| pAnt_*orf1*_R | tagtacGGATCCGGATCAAGGTTGCAATCG |
| pAnt_*orf2/3*_F | tagtacAAGCTTGACCGCGAGCACAAGTG |
| pAnt_*orf2/3*_R | tagtacGGATCCGGATCAGGAAAGCACGAAAC |
| ybbA_F | tagtacCTCGAGGCTGGCATTAACTACCGACG |
| ybbA_R | tagtacTCTAGAAGCCAGACAATTAATAGCGACG |

Restriction sites are highlighted in red. 6 random nucleotides were added at the 5'-end to improve DNA digestion with restriction enzymes (lowercase letters).
